# An ultrasound-guided biopsy technique for obtaining supraclavicular brown fat biopsies and preadipocytes

**DOI:** 10.1101/2024.04.30.570595

**Authors:** Eline S. Andersen, Naja Zenius Jespersen, Tora Ida Henriksen, Bente Klarlund Pedersen, Camilla Scheele, Michael Bachmann Nielsen, Søren Nielsen

## Abstract

Studying activated human brown adipose tissue (BAT) in vivo poses challenges due to its intricate anatomical positioning. Through the implementation of an ultrasound-guided biopsy technique, we successfully collected BAT samples from the supraclavicular region of 27 healthy individuals. As a comparative control, subcutaneous white adipose tissue (WAT) was similarly extracted from the same participants. Furthermore, we isolated progenitor cells from four tissue biopsies in both regions, subsequently subjecting them to a 12-day in vitro differentiation protocol following stimulation with 10 µM norepinephrine. To assess the mRNA expression of thermogenic genes within these small tissue samples, we employed a targeted cDNA amplification procedure, followed by conventional quantitative PCR (qPCR). Our study demonstrated that, with further refinement, this biopsy methodology can be used to obtain thermogenic adipose tissue. However, the expression data exhibited considerable diversity, and no statistically significant overall trends emerged for any of the five BAT marker genes (UCP1, PPARGC1A, PRDM16, CIDEA, CITED1), nor for the WAT marker HOXC8. The differentiation capacity of the progenitor cells revealed irregularities, with only three adipocyte cultures (two WAT and one BAT) displaying satisfactory differentiation potential. Remarkably, the differentiated BAT culture displayed a significantly elevated basal UCP mRNA expression level, further induced by 1.7-fold upon stimulation with norepinephrine. In summary, based on the in vitro data, brown adipose samples can be obtained using our ultrasound-guided biopsy technique approach. However, significant refinements are necessary before robust in vivo data can be generated in future intervention studies.

## Introduction

Since the discovery of brown adipose tissue (BAT) in adult humans[1–4], the tissue has been extensively investigated as a potential therapeutic target for obesity and its related metabolic complications[5–9]. Several studies have reported reduced volumes of BAT in people living with obesity or individuals with impaired cardiometabolic health [10,11]. Upon sympathetic activation, BAT consumes energy and takes up triglycerides and glucose, a feature that is utilized as the primary tool to quantify the volume and activity of human BAT depots[12]. The quantification of glucose uptake is used as an indirect measurement of volume, using a ^18^FDG glucose tracer for positron emission tomography coupled with computed tomography (FDG-PET/CT) scans. However, where the in vivo quantification in this fashion is a relatively uncomplicated procedure, studying the molecular events in the isolated tissue is far from trivial. The anatomical location of BAT in humans makes it difficult to obtain biopsies using conventional methods and involves several technical and practical challenges. Therefore, most of our knowledge about BAT biology is based on studies performed in rodents. However, the importance of performing in depth functional studies on brown fat in a species-specific context, was recently highlighted in a study showing that human BAT is activated by norepinephrine through β2-adrenergic receptors, and not β3-receptors as in rodents [13]. In addition, we have identified a human specific long noncoding RNA that regulates mitochondrial activity in BAT[14], a mechanism that is seemingly lacking in rodents. Thus, mapping and better understanding of the human molecular events in this tissue in humans is needed. In adults, the major BAT depots are located around the kidneys, along the spinal cord and in the deep neck and supraclavicular area[15–17], where especially the latter has been examined in various studies[18–21]. A common feature in these studies is that the adipose biopsies have been obtained during goitre or thyroid cancer surgeries where the neck region is already exposed[20] and while the patients were under general anaesthesia. This approach does not allow for in vivo intervention studies and as BAT is activated through the sympathetic nervous system, our knowledge about human brown adipose tissue is mainly limited to the inactivated state. For the same reason, some studies have utilised the Bergström needle method to take conventional biopsies in the supraclavicular region [22,23]. However, the supraclavicular region is a complex anatomical region which in addition to BAT comprises larger veins, arteries, nerve fibres and muscular tissues; therefore biopsies in that region are associated with significant risks[24]. A safer, less invasive, more reliable, and applicable biopsy method to study activated human BAT would thus be a major development within the research field. In this study, we therefore aimed to develop an ultrasound-guided small biopsy method to obtain brown adipose tissue depot from the neck region in healthy individuals. To validate the quality of the biopsies, we mapped the expression of thermogenic adipose tissue genes in BAT and used subcutaneous WAT as a reference. We furthermore isolated progenitor cells from both depots, which we expanded and differentiated in vitro. With a new method for obtaining human BAT, we would in the future be able to study activated BAT biology across different metabolic diseases and in response to different interventions such as cooling, exercise interventions and different postprandial states.

## Methods

### Study design

The study protocol was approved by the Scientific Ethical Committee of the capital region of Denmark, journal number H-1-2014-015, sub-study two. Healthy volunteers were recruited through advertisement. Healthy individuals between the age of 18 – 45 years with a BMI between 18 – 28 were eligible for participation. Furthermore, participants were required to have coagulation factors within the normal range i.e., INR: 0.9 – 1.2, APTT: 25 – 38 seconds, and thrombocytes > 145. Exclusion criterions were as follows: Use of daily medication besides anticonception medication for female participants, pregnancy, acute illness within two weeks prior to participation, chronic diseases including cancer, pulmonary-, cardiac-, kidney or liver-disease as well as endocrine disorders such as diabetes or thyroid disease, alcohol use of more than 14 units pr. week, or smoking in any form. Written informed consent was obtained from all participants following written and oral information about the study elements as well as anticipated risks and potential adverse events, and the study was conducted in accordance with the Helsinki declaration. Following an adverse event (see the results section) a modification to the protocol was included at the request of The Scientific Ethical Committee in which the study responsible doctor would contact the participants by telephone two days following the biopsy to ensure that there were no symptoms compatible with an adverse event.

All participants underwent a screening visit which consisted of a medical exam including an ECG, blood samples, the measuring of height, weight, waist- and hip circumference, a dual-energy X-ray (DXA) -scan, a two-hour oral glucose tolerance test, and a watt-max test.

Participants eligible for the biopsy visit met in the Hospital in the morning following 6 hours of fasting. Participants were instructed to use passive modes of transportation to get to the hospital and not to have performed vigorous exercise within 24 hours prior to the study day. An abdominal subcutaneous biopsy and one to two supraclavicular biopsies was performed as described in the biopsy section.

### Subjects

Thirty-two healthy volunteers (15 men and 17 women) with a median age of 25, range (18 – 44), BMI 24.5 (18.3 – 18.8), waist/hip ratio 0.8 (0.4 – 1.2) and total body fat percentage 28.6 (8.3 -50.1) were included in the study (table3, table 4). Two individuals were excluded following baseline examinations due to coagulation factors outside the normal range, and in three participants no biopsy could be obtained, due to insufficient fat in the region, evaluated by ultrasound (table 2). Two participants had two biopsies obtained on separate days, to be used for both tissue mRNA analyses and cell cultures. Thus, 58 biopsies were obtained from 27 subjects all together. 38 biopsies were snap-frozen in liquid nitrogen to be used for mRNA analyses, whereas 10 were utilized for pre-adipocyte isolation.

**Table 1.** The RNA yield from BAT and WAT lysates for each donor (anonymized ID). Nine samples had RNA concentration below the Qubit detection threshold (N.D.).

| Tissue ID (Anonymized) |  | RNA Yield (ng/ul) |  | qPCR signal (5ng cDNA) |  |
| --- | --- | --- | --- | --- | --- |
| BAT | WAT | BAT | WAT | BAT | WAT |
| 1B | 1W | N.D. | 2.31 | + | + |
| 2B | 2W | N.D. | N.D. | + | + |
| 3B | 3W | 1.6 | 1.1 | + | + |
| 4B | 4W | 1.97 | 1.29 | + | + |
| 5B | 5S | 1.36 | 1.46 | + | + |
| 6B | 6W | N.D. | 1.47 | + | + |
| 7B | 7W | N.D. | 1.58 | + | + |
| 8B | 8W | 2.1 | N.D. | + | + |
| 9B | 9W | 1.65 | 3.5 | + | + |
| 10B | 10W | 12.2 | 10 | + | + |
| 11B | 11W | 2.43 | 9.85 | + | + |
| 12B | 12W | N.D. | 6.2 | + | + |
| 13B | 13W | 1.81 | 3.9 | + | + |
| 14B | 14W | 15 | 3.98 | + | + |
| 15B | 15W | 2.34 | 4.05 | + | + |
| 16B | 16W | N.D. | 2.76 | % | + |
| 17B | 17W | N.D. | 2.16 | + | + |
| 18B | 18W | 1.41 | 2.72 | + | + |
| 19B | 19W | 1.5 | 3.78 | + | + |

**Table 2.** Inclusion and exclusion criterions for participation in the study.

|  |  |
| --- | --- |
| Inclusion Criteria | - Age 18 – 45 years<br>- Body mass index (BMI) 18 – 29 kg/m<sup>2</sup><br>- Coagulation factors within the normal range i.e. platelet count > 145, International Normalized ratio (INR) 0.9 -1.2, and activated partial thromboplastin time (APTT) 25 – 38 seconds. |
| Exclusion criteria | - Use of daily medications, apart from oral contraceptives for the women.<br>- Acute illness within two weeks prior to the experimental day<br>- Chronical illnesses, including cancers, metabolic diseases, such as diabetes or thyroid disease, and chronic heart- kidney-, liver-, and pulmonary diseases<br>- Alcohol consumption of more than 14 units / week<br>- Smoking<br>- Pregnancy for the women |

**Table 3.** Participant characteristics, tissue samples. Values are median and range. F = Female, M = male, BMI = Body Mass Index, WHR = Waist Hip ratio.

| Sex | Age (years) | BMI (kg/m<sup>2</sup>) | WHR | Fat percentage |
| --- | --- | --- | --- | --- |
| F (11) | 24.5 (20.7 – 36) | 23.5 (18.6 – 28.8) | 0.79 (0.69 – 1.21) | 36.5 (22.5 – 50.1) |
| M (9) | 24.8 (19.3 -44) | 26.2 (22.5 – 28.4) | 0.97 (0.82 – 1.02) | 20.5 (8.5 -33.4) |

**Table 4.** Participant characteristics, Cell culture samples. Values are median and range. F = Female, M = male, BMI = Body Mass Index, WHR = Waist Hip ratio.

| Sex | Age (years) | BMI (kg/m<sup>2</sup>) | WHR | Fat percentage |
| --- | --- | --- | --- | --- |
| F (4) | 26.3 (23.2 – 31) | 24.4 (21.8 – 24.6) | 0.68 (0.39 – 0.79) | 19.25 (8.5 – 50.1) |
| M (6) | 25.0 (19.3 -35.8) | 25.2 (22.2 – 26.7) | 0.88 (0.82 – 0.79) | 36.9 (33.6 - 38.4) |

All included subjects had normal glucose tolerance, as well as TSH, HBA1c, high sensitive CRP, white blood cell count, hemoglobin and cholesterol, including both HDL, LDL and triglyceride levels within the normal range.

### Biopsy

All participants were initially examined by ultrasound B-mode to localize the adipose tissue. A GE Logiq E9 system (GE Healthcare, Chalfont St. Giles, UK) with a linear array, high frequency probe (ML6-15) was used for all examinations. Participants were supine with the neck extended, exposing the supraclavicular area. Both sides of the neck were examined and the side with the easiest and safest access for biopsy was chosen. Local anaesthesia (2 ml of Lidocaine 20mg/ml) was injected subcutaneously. Afterwards the skin was sterilized with ethanol and a small incision was made in the skin. Ultrasound guided biopsy was performed two to three times with a semi-automatic 18G biopsy needle (Medax, Mantova, Italy). The same experienced radiologist (>20 years of interventional ultrasound) performed all supraclavicular biopsies. The biopsy was cleansed using sterile saline and either immediately snap-frozen in liquid nitrogen or placed in cell culture medium (DMEM / F12) + 1% PS. Afterwards an abdominal control biopsy was obtained from the periumbilical subcutaneous adipose depot by one the study investigators using the same method but without ultrasound guidance.

### RNA isolation

The RNeasy Micro Kit (Qiagen) was used to isolate the RNA from the small biopsies. The kit contains poly-A RNA for use as carrier RNA to improve the isolation efficiency from very small tissue samples. The carrier RNA approach was tested according to the manufacturer’s guidelines. The isolation efficiency was compared to a control with no poly-A carrier added. As there was no difference in efficiencies and a potential risk for ineffective amplification with carrier RNA added, we proceeded with the isolation procedure without the carrier. A lysis buffer consisting of 10 ul β-Mercaptoethanol (Sigma, 63689-25ML-F) per 1 ml RLT Buffer (included in the RNeasy Micro Kit) was prepared. The tissue was placed in 2 ml microcentrifuge tubes containing one 5 mm stainless steel bead and added 350 µl lysis buffer per tube. Tubes was placed in the TissueLyser and the tissues were homogenized for 2 min at 20 Hz. Tubes were then transferred to a centrifuge and centrifuged at 16,000 rpm for 3 min and 30 µl of the supernatant was then transferred to new 2 ml tubes containing 350 ul 70% ethanol and mixed with a pipette. Samples were then transferred to RNeasy MinElute spin column placed in a 2 ml collection tube and then centrifuged for 15 s at 10,000 rpm. Columns were then washed with 350 µl RW1 buffer (included in the RNeasy Micro Kit) and then incubated with an 80 µl DNAse buffer (1500 Kunitz unit DNAse solution in 70 µl RDD Buffer) for 30 min and then washed with provided buffers and 80% ethanol, according to the manufacturer’s guideline. Lastly, 14 µl RNasefree water (Thermo Fisher) was added to the samples to elute RNA from the spinning filters. RNA concentrations were determined with an RNA High sensitivity assay (Thermo Fisher) using a Qubit 3.0. (table 1)

### cDNA procedure and qPCR

#### Adipose Tissue

Total RNA (5 ng) was reverse transcribed using cDNA high-capacity kit (Applied Biosystems) according to the manufacturer’s protocol. For those 9 samples (table 1) that were out of detection range, a 5 µl lysate volume was used as input. Then the cDNA was preamplified using the Taqman PreAmp Master Mix (Applied Biosystems) targeting 33 defined mRNAs (Supplementary table 1). Briefly, 5 ul of 33 Taqman assays (Supplementary table 1) was pooled in a tube and 6,25 µl of the taqman pool was then mixed with 6,25 µl cDNA and 12.5 preamp master mix. Tubes were placed in a Thermo cycler and preamplified at 10 cycles (denature 95°C for 15 sec, 60°C for 4 min). Diluted cDNA samples (1:10) were loaded in duplicates and qPCR was performed using ViiA7 Sequence Detection system (Applied Biosystems, Foster City, CA, USA) according to the manufacturer’s protocol using TaqMan™ Universal PCR Master Mix (Thermo Fisher).

#### Primary adipocyte cultures

Total RNA (200 ng) was reverse transcribed using a cDNA High-Capacity Kit (Applied Biosystems) according to the manufacturer’s protocol. cDNA samples were loaded in triplicate and qPCR was performed with the ViiA7 Sequence Detection system (Applied Biosystems), according to the manufacturer’s protocol, using PowerUp SYBR Green Master Mix for quantification of *FABP4, PPARGCA, PPIA* and *PRDM16* mRNA (Primer sequences below). *UCP1* mRNA was quantified using TaqMan Universal PCR Master Mix and the TaqMan primer/probe assay Hs00222453_m1 (ThermoFischer Scientific). SYBR-based primers were designed using the Roche Applied Science Assay Design Centre (Roche) and checked for specificity using Primer-Blast (NCBI).

Primers:

FABP4 Forward: CCTTTAAAAATACTGAGATTTCCTTCA

FABP4 Reverse: GGACACCCCCATCTAAGGTT

PPARGC1A Forward: CAAGCCAAACCAACAACTTTATCTCT

PPARGC1A Reverse: CACACTTAAGGTGCGTTCAATAGTC

PPIA Forward: ACGCCACCGCCGAGGAAAAC

PPIA Reverse: TGCAAACAGCTCAAAGGAGACGC

PRDM16 Forward: CACGAGTGCAAGGACTGC

PRDM16 Reverse: TGTGGATGACCATGTGCTG

### Cell culture of human primary adipocytes

Preadipocytes were isolated from the small supraclavicular and abdominal subcutaneous adipose tissue biopsies. A previously published isolation protocol was modified to preserve as many viable preadipocytes from the very small biopsies as possible material. The adipocyte differentiation procedure was as previously published[20,25]. Briefly, biopsies were digested in DMEM/F12 containing collagenase II (1 mg/ml; Sigma Aldrich) and fatty acid-free bovine serum albumin (15 mg/ml; Sigma Aldrich). The solution was passed through a cell strainer which was subsequently flushed with 10 ml DMEM/F12 to capture as many cells as possible. The solution was centrifuged at 800g for 7 minutes, and cell pellet was subsequently washed with DMEM/F12. The cell pellet was resuspended in growth media consisting of DMEM/F12, 10% Fetal bovine serum, 1% Penicillin-Streptomycin and cells were seeded in one well in a 12-well plate. The cells were grown at 37°C in an atmosphere of 5% CO2 and the medium was changed every second day. Adipocyte differentiation was induced two days after preadipocyte cultures were 100% confluent by treating cells with DMEM/F12 containing 1% Penicillin-Streptomycin, 0.1 μM dexamethasone (Sigma-Aldrich), 100 nM insulin (Actrapid, Novo Nordisk or Humulin, Eli Lilly), 200 nM rosiglitazone (Sigma-Aldrich), 540 μM isobutylmethylxanthine (IBMX) (Sigma-Aldrich), 2 nM T3 (Sigma-Aldrich) and 10 μg/ml transferrin (Sigma-Aldrich). After three days of differentiation, IBMX was removed from the cell culture media. The cell cultures were left to differentiate for an additional nine days, with medium change the third day. Following 12 days of differentiation, cells were harvested for RNA, protein. When stated in the figure legend, cells were stimulated with 10 μM norepinephrine (Sigma-Aldrich) for 4 hr before RNA and protein were isolated. Two hours prior to the norepinephrine stimulation, old medium was replaced by DMEM/F12 (Life technologies) containing 1% penicillin-streptomycin.

### Statistics and data availability

The statistical analysis was performed using GraphPad Prism 9.0 (GraphPad Software Inc., La Jolla, CA, USA). Results are presented as means ± S.E.M. Tissue qPCR data did not pass the Kolmogorov–Smirnov test for normal distribution and therefore data were analysed using the nonparametric Wilcoxon matched-pairs signed rank test. Cell qPCR data were analysed using a 2-way ANOVA. All raw PCR are available online at Mendeley Data.

## Results

### An ultrasound guided small needle approach

Using an ultrasound guided small biopsy needle approach we aimed for obtaining sufficient volume of adipose tissue to 1) extract RNA that could be further evaluated with qPCR and 2) isolation of adipose progenitors that could be expanded and differentiated to mature adipocytes (Figure 1A).

**Fig. 1.**
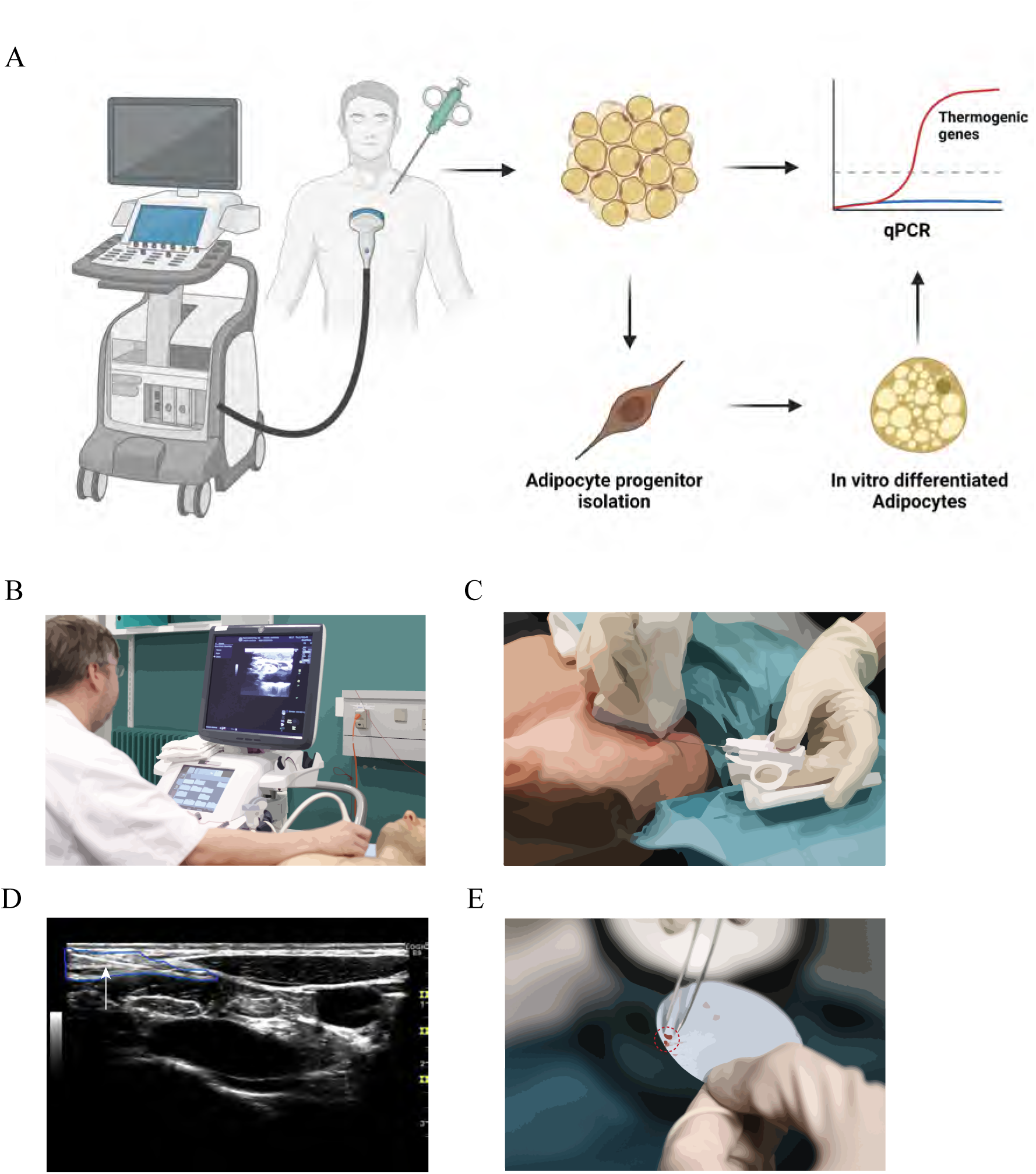
Ultrasound guided biopsy method for obtaining brown adipose tissue in humans. **A,** Graphical overview on the experimental design. **B**, Ultrasound guided localisation of the supraclavicular region. **C**, Small needle insertion (semi-automatic 18G biopsy needle) in the supraclavicular neck area for obtaining brown adipose tissue. **D**, Image of the supraclavicular neck area generated by ultrasound. The biopsy needle is outlined with blue. E, Example of the adipose tissue size after the biopsy procedure.

All participants were initially examined, in a supine position, exposing the supraclavicular neck area by ultrasound to localize regions with adipose tissue (figure 1B, 1C). Ultrasound-guided biopsy was performed two to three times with a semi-automatic 18G biopsy needle while carefully monitoring the needle position (Figure 1D). Small adipose biopsies were successfully taken out from the supraclavicular region for further examination (Figure 1E). Subcutaneous abdominal adipose tissue was used as reference material using the same biopsy method.

### Applicability and adverse events

The biopsy method was fast and easily applicable, and aside from a slight burning sensation related to the injection of local analgesia most subjects felt no discomfort in relation to the procedure. This was with the except of one case, in which the supraclavicular procedure itself caused an immediate sharp pain. In the following days, the subject developed an iatrogenic pneumothorax which gradually increased in size over the next week to require drainage. Thus, the subject had to be hospitalized and required a chest drain for 5 days. On this treatment the pneumothorax was resolved, and the subject recovered without any permanent sequelae. There were no other cases of adverse events or complications.

### Evaluation of depot specific mRNA signature in small adipose tissue biopsies

Based on the *PPIA* expression, we succeeded in isolating sufficient concentrations of RNA from all the biopsies except from 1 sample (table 1). Furthermore, as expected for a housekeeping gene, the expression levels were comparable between WAT and BAT depots and expressed at levels comparable with higher RNA input approaches (WAT: mean CT=27.4 +/-1.2 SD, BAT: CT=28.1 +/- 1.3 SD) (Figure 2B). We next evaluated the expression of the well-described markers of brown and white adipose identity *UCP1*, *PPARGCA*, *CITED1*, *PRDM16*, *CIDEA* (all BAT) and *HOXC8* (WAT). Apart from *CIDEA* and *HOXC8*, the expression of the identity markers was either lowly expressed or not detectable after 45 qPCR cycles. When comparing the expression levels between the two depots, *CIDEA* mRNA expression was surprisingly higher in WAT compared to BAT (p<0.01) (Figure 2C). None of the other measured markers were expressed different between the two depots (figure 2C).

**Fig. 2.**
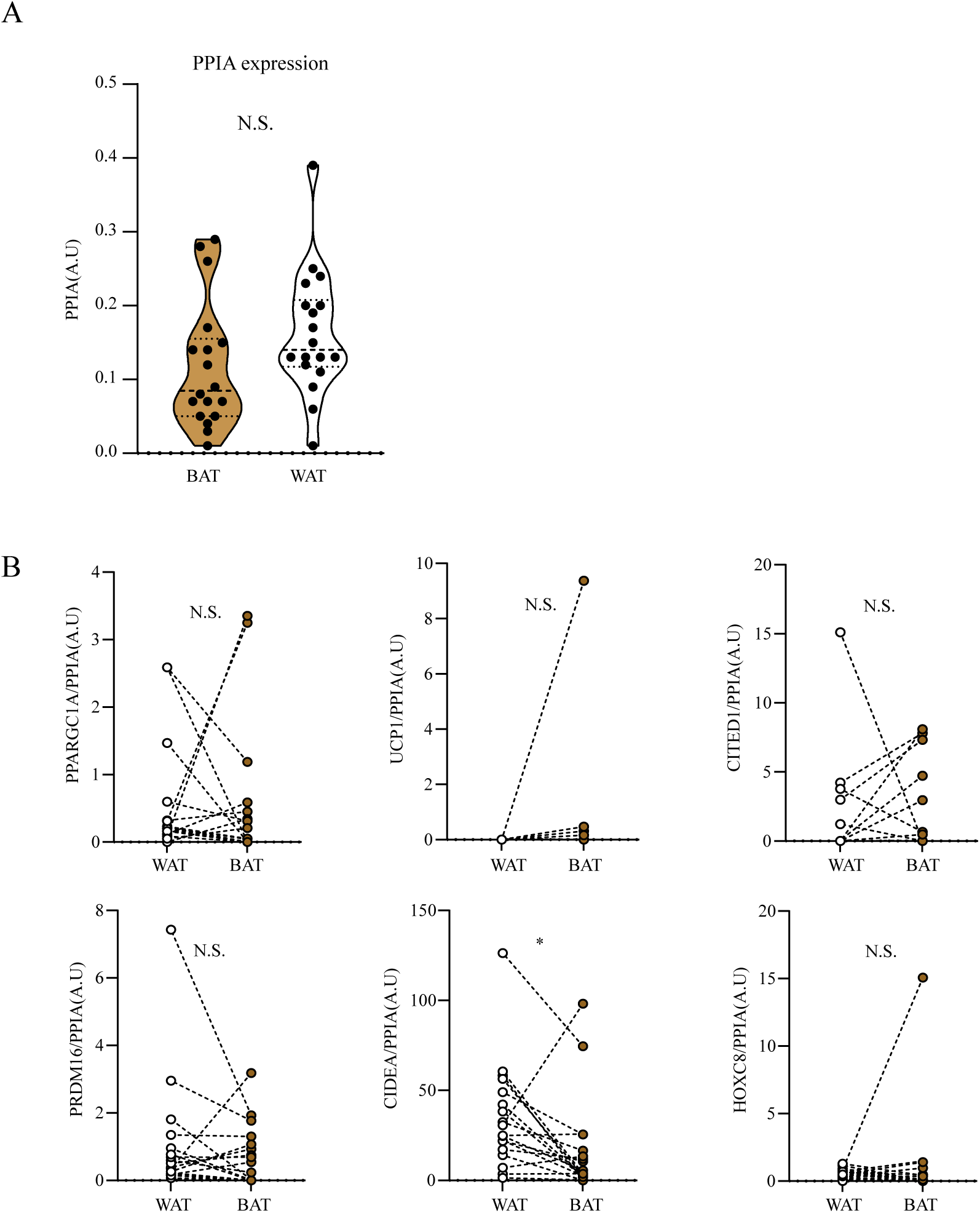
Evaluation of thermogenic mRNA markers in adipose tissue biopsies obtained. **A**, Evaluation of cDNA efficiency based on the PPIA mRNA expression (n=19). **B**, Evaluation of the expression of thermogenic mRNA markers (UCP1, CITED1, PPARGC1A, CIDEA) and the white adipose tissue marker (HOXC8). Statistical significance was determined by Wilcoxon matched pairs signed rank test (n=18) * = p<0.05.

### Evaluation of adipogenic capacity and thermogenic potential in vitro

Out of the ten paired cultures, we succeeded isolating progenitors from 4 WAT and 4 BAT depots (not paired). We conclude that a higher volume of adipose tissue, or possibly further refinement of the isolation protocol is needed to secure expansion and differentiation of the progenitor cells. Importantly, we have later shown that expansion of already isolated progenitorsfrom single cells is possible when provided conditioned media from growing adipose progenitors (REF to Nat Met paper). Thus, using a 48-well plate or even a 96-well plate and providing conditioned media, would potentially result in securing cell isolation from small-sized biopsies in the future.

All eight cultures could be expanded until 100% confluence, followed by initiation of the adipocyte differentiation program. However, only three cultures (Two WAT cultures and one BAT cultures) had the potential to differentiate into mature adipocytes, based on the cellular lipid accumulation (figure 3A-B). We have recently described that cells divide into two major cell fates during differentiation: adipogenic and SWAT cells[26] . Whereas the adipogenic cells accumulate lipids, the SWAT cells express extracellular matrix factors. Many of the current cultures thus appears to be dominated by SWAT cells. In terms of thermogenic capacity, there was no statistically significant difference in thermogenic gene expression between depots or in response to a stimulus with norepinephrine (Figure 3C). However, expression of *UCP1* and *PPARGC1A* mRNA could be induced in individual cultures. We have previous experience in that there is a large variation in terms of lipid droplet accumulation as well as thermogenic capacity[19]. Therefore, we cannot conclude on whether that the current results depend on technical limitations of the small biopsies or individual variation in thermogenic potential.

**Fig. 3.**
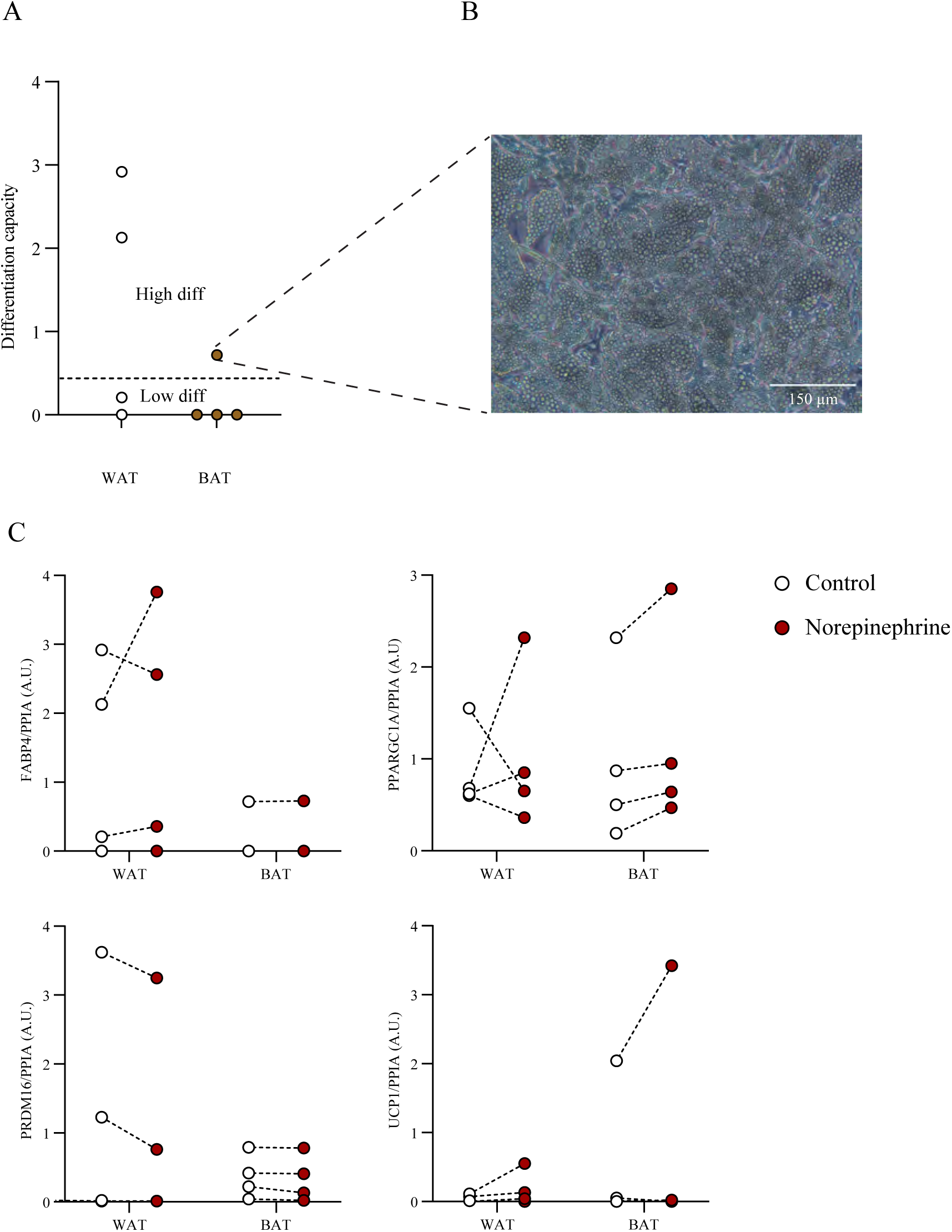
Evaluation of thermogenic mRNA markers in in vitro differentiated progenitors obtained with ultrasound guided small biopsy needle technique. **A,** Differentiation capacity based on FABP4 mRNA expression (n=4). **B** Brightfield Image of fully differentiated adipocytes obtained with small needle biopsy technique.). **C** Thermogenic mRNA marker expression (PRDM16, PPARGC1A, UCP1, FABPF) in brown and white adipocytes after 12-day differentiation period, stimulated with 10 μM norepinephrine for 4 hours

## Discussion

In this project we developed a cost-effective method for obtaining samples of human supraclavicular adipose tissue and evaluating the thermogenic mRNA signature as well as isolating preadipocytes from the sample. However, several technical and practical challenges need to be solved before the method can be implemented.

In this project we assumed that all participants had brown adipose tissue in the supraclavicular region, which we could reach with the biopsy needle and further process for analysis. However, the BAT volume in humans is very variable and are for some individuals completely absent[20]. For instance, studies have shown a dramatic decrease of BAT prevalence with age[10]. Furthermore, there are several reports BAT volume being correlated with BMI, glucose tolerance, gender, and ethnicity[10,27]. Therefore, a uniform cohort of younger people with known BAT depots of significant volume should be selected as an inclusion criterion. Although the participants in the present study were relatively young, healthy, and non-obese, a pre-screening procedure including a PET-CT scan combined with a cooling protocol [28] to ensure the presence of BAT depots of significant volume should be included. A detailed PET-CT BAT localisation map for each individual would most likely improve the BAT biopsy accuracy.

As described in the introduction, there are several risks associated with performing biopsies in the supraclavicular region including the risk of bleeding, infection, nerve damage and development of a pneumothorax. In a previous study, where a Bergström needle was utilized 9.6% of participants developed transient numbness and 2.3% hematoma[23]. In the present study, three subjects were excluded due to insufficient amounts of supraclavicular fat for the biopsy to be performed in a safe manner. In spite of this, one subject nevertheless developed a serious complication in the form of a pneumothorax which required hospitalization and drainage. We assume that the utilization of a smaller needle gauge is the underlying factor contributing to the absence of any additional complications in the current study. Importantly, the observed pneumothorax underlines the difficulty in the application of the described biopsy method.

In addition, alternatives in tissue handling and the wet lab procedures needs to be considered. First, a gentler tissue lysis procedure could improve the RNA yield. In this project, we used a silver bullet approach that ruptures the tissue in 2 ml Eppendorf tubes. This method is very efficient when processing larger tissue samples. However, as the tissue in this project was barely visible, it was difficult to evaluate the lysis efficiency. A method with smaller volume of lysis buffer and a manual lysis procedure with a pipette might have resulted in higher RNA yield. Furthermore, the cDNA concentration was in general low and the selected targeted preamplification approach might not be the most efficient method for this purpose. Alternative and more advanced protocols within the single cell technology have been in rapid development within the recent years[29,30] . These technologies are today available as commercialized kits and would most likely have been beneficial for analysing the material isolated in this project.

While the biopsy procedure and the processing of the tissue for mRNA-analysis could be improved, the volume of the tissue may be too low for successful isolation of adipogenic progenitors. However, the protocol for cell isolation used in this study[20] was a modified protocol optimized for larger biopsies and could be revised even further for better performance on smaller biopsies. For instance, a protocol allowing to isolate and culture skin cells from a single 4 mm-punch biopsy, has been developed[31]. The sample and cell size corresponds to the material this method could most likely be applied in a future protocol.

Cell death inducing DFFA like effector a (CIDEA) is a well-established brown adipose tissue mRNA marker[32]. However, in this study we show an unexpected higher expression of *CIDEA* in the subcutaneous white adipose tissue. Despite reports about the importance of CIDEA in white adipose tissue[33], the finding is more likely connected to a technical artefact.

In conclusion, we present an ultrasound guided method for obtaining biopsies from the human supraclavicular adipose depot. We show that preadipocytes can be isolated from the samples and that it is possible to extract sufficient RNA for qPCR from the samples. The method, however, needs further development to improve the success rate.

With a new and improved method for obtaining human BAT, we would in the future be able to study activated BAT biology across different metabolic diseases and in response to different interventions such as cooling, exercise interventions and different postprandial states.

## Acknowledgement

We acknowledge Noemi James, Lone Peijs and Maria Scheel for the technical assistance during the study period. Furthermore, we acknowledge all the included voluntary study participants for their time, trust, and curiosity.

