## Supplementary material for "An ultrasound-guided biopsy technique for obtaining supraclavicular brown fat biopsies and preadipocytes": Suplementary table 1

| Gene list | Assay | Dye |
| --- | --- | --- |
| ZIC1 | Hs00602749_m1 | FAM-MGB |
| TBX1 | Hs00962556_m1 | FAM-MGB |
| CIDEA | Hs00154455_m1 | FAM-MGB |
| CITED 1 | Hs00918445_g1 | FAM-MGB |
| TMEM26 | Hs00415619_m1 | FAM-MGB |
| HOXC9 | Hs00396786_m1 | FAM-MGB |
| EBF3 | Hs00406051_m1 | FAM-MGB |
| FBXO31 | Hs00375551_m1 | FAM-MGB |
| EPSTI1 | Hs01566789_m1 | FAM-MGB |
| CD137 | Hs00155512_m1 | FAM-MGB |
| RXRg | Hs00199455_m1 | FAM-MGB |
| Myf5 | Hs00929416_g1 | FAM-MGB |
| FGF21 | Hs00173927_m1 | FAM-MGB |
| BMP4 | Hs00370078_m1 | FAM-MGB |
| BMP7 | Hs00233476_m1 | FAM-MGB |
| DIO2 | Hs00988260_m1 | FAM-MGB |
| ADRb3 | Hs00609046_m1 | FAM-MGB |
| KCNK3 | Hs00605529_m1 | FAM-MGB |
| HMGCS2 | Hs00985427_m1 | FAM-MGB |
| CKMT2 | Hs00176502_m1 | FAM-MGB |
| TGM2 | Hs00190278_m1 | FAM-MGB |
| ARG2 | Hs00982833_m1 | FAM-MGB |
| COBL | Hs00391205_m1 | FAM-MGB |
| NTRK3 | Hs00176797_m1 | FAM-MGB |
| LEP /leptin ( | Hs00174877_m1 | FAM-MGB |
| TCF21 | Hs00162646_m1 | FAM-MGB |
| UCP1 | Hs00222453_m1 | FAM-MGB |
| HOXC8 | Hs00224073_m1 | FAM-MGB |
| PPIA | Hs01565699_g1 | FAM-MGB |
| PGC-1(alpha) | Hs01016724_m1 | FAM-MGB |
| PRDM16 | Hs00922674_m1 | FAM-MGB |
| FABP4 | Hs01086177_m1 | FAM-MGB |
| LHX8 | Hs00418293_m1 | FAM-MGB |
